# Noradrenergic alpha-2a Receptor Stimulation Enhances Prediction Error Signaling in Anterior Cingulate Cortex and Striatum

**DOI:** 10.1101/2023.10.25.564052

**Authors:** Seyed A. Hassani, Thilo Womelsdorf

## Abstract

The noradrenergic system is implicated to support behavioral flexibility by increasing exploration during periods of uncertainty and by enhancing working memory for goal-relevant stimuli. Possible sources mediating these pro-cognitive effects are α2A adrenoceptors (α2AR) in prefrontal cortex or the anterior cingulate cortex facilitating fronto-striatal learning processes. We tested this hypothesis by selectively stimulating α2ARs using Guanfacine during feature-based attentional set shifting in nonhuman primates. We found that α2A stimulation improved learning from errors and facilitates updating the target feature of an attentional set. Neural recordings in the anterior cingulate cortex (ACC), the dorsolateral prefrontal cortex (dlPFC), and the striatum showed that α2A stimulation selectively enhanced the neural representation of negative reward prediction errors in neurons of the ACC and of positive prediction errors in the striatum, but not in dlPFC. This modulation was accompanied by enhanced encoding of the feature and location of the attended target across the fronto-striatal network. Enhanced learning was paralleled by enhanced encoding of outcomes in putative fast-spiking interneurons in the ACC, dlPFC, and striatum but not in broad spiking cells, pointing to an interneuron mediated mechanism of α2AR action. These results illustrate that α2A receptors causally support the noradrenergic enhancement of updating attention sets through an enhancement of prediction error signaling in the ACC and the striatum.

## Introduction

The noradrenergic system supports attention, working memory and learning functions (Berridge and Waterhouse, 2003; Aston-Jones and Cohen, 2005; Bouret and Sara, 2005; Doya, 2008; Sara, 2009; Silvetti et al., 2013; Bouret and Richmond, 2015; Poe et al., 2020; Bornert and Bouret, 2021; Holland et al., 2021). These higher cognitive functions are critically supported by bi-directional projections of noradrenergic neurons in the locus coeruleus and neurons in the anterior cingulate and prefrontal cortex (Arnsten and Goldman-Rakic, 1984; Tervo et al., 2014; Cope et al., 2019; Bari et al., 2020; Su and Cohen, 2022). Optogenetic evidence has directly implicated noradrenergic activation to support adaptative behaviors. Noradrenergic neurons in the locus coeruleus projecting to the frontal cortex of mice show higher firing rates when a choice leads to an unexpected outcome that requires adjusting behavior (Su and Cohen, 2022). Silencing the noradrenergic projections to the frontal cortex in mice impairs behavioral switching (Su and Cohen, 2022), while enhancing noradrenergic neuron activation in the locus coeruleus can increase the speed of behavioral switching in mice (McBurney-Lin et al., 2022). These recent insights suggest that noradrenergic modulation of prefrontal circuits might be a key mechanism in the nonhuman primate (NHP) brain for the flexible switching of complex cognitive states such as the updating of attention sets.

There are various ways how noradrenergic modulation could support the flexible switching of cognitive states. One possibility is that tonic noradrenergic enhancement modulates arousal (Rajkowski et al., 1994), and increases exploratory behaviors (Koralek and Costa, 2021). However, while noradrenergic modulation can have positive effects by ‘releasing’ behavior from following expected values in order to seek novel and potentially more valuable information (Jahn et al., 2023), stimulation of noradrenergic neurons may also shift behavior to randomness (Tervo et al., 2014) or trigger behavioral shifts away from still valuable foraging patches, reflecting maladaptive or suboptimal behaviors (Kane et al., 2017). Another means for noradrenaline to support the flexible switching of cognitive states is by enhancing the signal-to-noise ratio (SNR) of neuronally encoded salient events including enhanced representations of task relevant stimuli (Kolta et al., 1987; Kossl and Vater, 1989; Waterhouse et al., 1990; Ciombor et al., 1999; Devilbiss and Waterhouse, 2004; Decamp et al., 2011; Ghosh and Maunsell, 2022). For example, noradrenergic stimulation of the lateral prefrontal cortex can enhance persistent delay firing of neurons when they encode their preferred spatial location in short-term memory (Wang et al., 2007). However, persistent working memory representations in lateral prefrontal cortex cannot easily explain cognitively flexible behavior, which is characterized by rapidly changing working memory representations to varying task demands and reward contingencies (McDougle and Collins, 2021; Womelsdorf et al., 2021), which are processes more closely associated with medial and orbital prefrontal circuits rather than with lateral prefrontal cortex (Sadacca et al., 2017).

In order to shed light on how noradrenergic modulation improves cognitive flexibility and neural signaling we set out testing whether enhancing α2A noradrenergic receptors is sufficient to improve cognitive flexibility and enhance neuronal encoding of learning-relevant task variables in the anterior cingulate cortex (ACC), the dorsolateral prefrontal cortex (dlPFC, areas 9/46d and 9/46v), and the head of the caudate nucleus (striatum). We investigated these questions by administering the α2A receptor specific drug Guanfacine in NHP performing an attention set shifting task while we recorded single neuron activity in ACC, dlPFC, and striatum. NHP lesion studies have documented these areas are necessary for adaptive learning and each area contains neurons with activity correlating with efficient learning of abstract feature values during set shifting (Kennerley et al., 2006; Buckley et al., 2009; Heilbronner et al., 2011; Gläscher et al., 2012; Oemisch et al., 2019; Boroujeni et al., 2020, 2021; Passingham, 2021). The task reversed color-reward associations of an attention set (**Figure 1A**) and dissociated covert feature-based attention to color from spatial attention to peripheral stimulus locations and from the motor requirements to make a directional saccade (**Figure 1B**). We chose Guanfacine because (*i*) it is a highly selective α2A receptor agonist with 15-60x higher affinity for the α2A, over the α2B and α2C subtypes (Uhlen and Wikberg, 1991; Uhlen et al., 1994), (*ii*) its mechanism enhancing working memory in neurons of the dlPFC is well documented (Arnsten and Goldman-Rakic, 1985; Li and Mei, 1994; Arnsten et al., 1996; Mao et al., 1999; Arnsten, 2000; Wang et al., 2007), while (*iii*) its possible modulation of flexibility has remained unclear (Berridge and Spencer, 2016).

**Figure 1.**
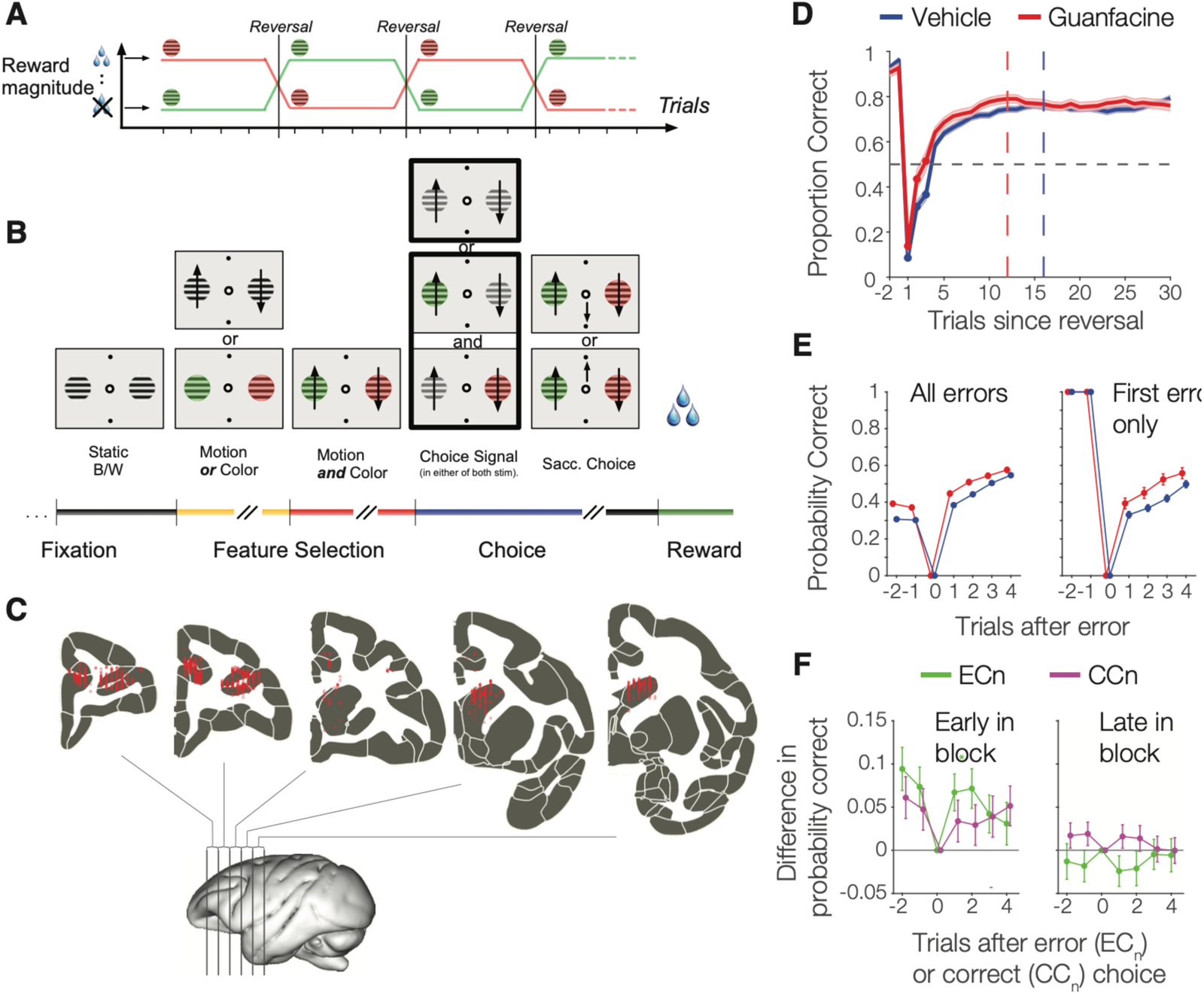
Task paradigm anatomical recording locations and behavioral results. (**A**) Monkeys performed a feature-based reversal learning task that rewarded one of two colors in blocks, reversing the rewarded color without cue when at least 30 trials were completed and a 90% performance criterion (over preceding 12 trials) was reached. (**B**) In each trials monkeys fixated and covertly processed color and a motion onset of two peripheral stimuli. One, or both stimuli transiently dimmed at an unpredictable time and subject had to ignore the direction of motion of the distractor, while making a saccade in the direction of the target stimuli. (**C**) Reconstructed electrode recording locations in dlPFC, ACC and striatum. (**D**) Performance for learned blocks (first block excluded) in Guanfacine and vehicle sessions (shading denote SE). Learning curves were smoothed (4-trial window) from trial 4 onwards. Horizontal dashed line shows chance probability. Vertical dashed lines represent the median first trial at which criterion proportion correct responses of p≥0.7 (over successive 10 trials) was reached for Guanfacine (12) and vehicle (16). (**E**) Proportion correct in trials before and after error trials (EC_n_ analysis) with Guanfacine (red) and vehicle (blue). Right panel: proportion correct after the first error in a block. (**F**) EC_n_ and CC_n_ (proportion correct after correct trials) analysis for the first 9 trials in a block (*left*) and for trials 10 onwards (*right*).

We found that α2A stimulation improves using error feedback to adjust behavior and speeds up reversal learning of attention sets. These behavioral improvements were paralleled by enhanced encoding of negative prediction errors in ACC and of positive prediction errors in the striatum. These findings illustrate that the α2A receptor system is sufficient to enhance flexible reversal learning of attention sets and they identify the ACC – striatum circuit as a system translating α2A stimulation into improved behavior.

## Results

Two rhesus monkeys performed the attention set shifting task for 42 (monkey H, 17 with Guanfacine injections) and 66 (monkey K, 12 with Guanfacine injections) electrophysiological recording sessions respectively. A total of 1157 single units (619 and 538 units from monkey H and K) were collected across the ACC (area 24, n = 456 units), dlPFC (areas 8a/46d/46v, n = 410 units), and in the anterior striatum (n = 291 units) (**Figure 1C**).

### Guanfacine enhances reversal learning and post-error adjustment

We quantified the speed of reversal learning on days with vehicle and Guanfacine administration as the first trial in a block at which performance exceeded the learning criterion (p=.70 correct over subsequent 10 trials). Average learning occurred at trial 16 and occurred 3.2 trials earlier with Guanfacine than with vehicle (p = .003; **Figure 1D**), which was evident in each monkey (**Figure S1A**) and when estimating learning speed with an ideal observer statistics (**Figure S1B**). Faster learning depended on reversing a previously learned color-reward association as it started with the first reversal block of a session (**Figure S1C**). Guanfacine did not alter the overall proportion of reversal blocks that were learned (n.s.).

Faster learning could be achieved by adjusting performance after nonrewarded (error) trials or after rewarded (correct) choices. We tested for changes in performance accuracy post-error and post-correct trials early or later in a block and found a significant main effect of drug condition (F(1,412) = 4.89; p = .028), block timing (F(1,412) = 62.54; p < .001) and post-error vs post-correct trials (F(1,412) = 266.48; p < .001) with a significant interaction between drug condition and timing in the block (F(1,412) = 6.63; p = .010). These results reflect that early in the block (first 9 trials), Guanfacine significantly improved post-error performance for 2 trials after an error (6.9%; p = .022) (**Figure 1E**), but did not alter post-correct performance after rewarded trials (3.2%; p = .809) (**Figure 1F**). The post-error enhancement of performance with Guanfacine was already observable after the very first error trial in a new reversal block (**Figure 1E** *right*). Later in the block (from trials 10 onwards), Guanfacine did not affect performance after error (2.3%; p = .963) or correct trials (1.5%; p = .997).

### Pupils constrict with Guanfacine

We analyzed the pupil diameter of the monkeys to infer the effect of Guanfacine on noradrenergic neuron activity in the locus coeruleus (LC). During the first three blocks of each session, when Guanfacine concentrations are highest, monkeys had a more constricted pupil diameter compared to vehicle (monkey H: p < .001; monkey K: p = .010; **Figure S1D**). This finding suggests Guanfacine reduced LC activity as expected from physiological studies that showed a2A-noradrenergic receptors are expressed pre-synaptically on LC terminals where they act as inhibitory auto-receptors that decrease noradrenergic release (Engberg and Eriksson, 1991; Okada et al., 2019).

### Guanfacine does not alter overall firing rates or firing variability

Guanfacine’s behavioral effects might relate to overall firing rate changes in ACC or dlPFC based on studies indicating Guanfacine increases excitability in prefrontal pyramidal neurons (Wang et al., 2007; Barth et al., 2008; Kawaura et al., 2014). Since α2A adrenoceptors are likely differentially expressed in pyramidal neurons and interneurons (Kawaguchi and Shindou, 1998; Wang et al., 2013; Liu et al., 2014; Xing et al., 2016; Lee et al., 2020) we first split the neurons into narrow and broad spiking (NS and BS) neurons. Previous work has shown that fast spiking interneurons have faster (narrower) spike-width (Ardid et al., 2015; Oemisch et al., 2019), and that neurons with narrower spike-width suppress multiunit activity consistent with being inhibitory (Boroujeni et al., 2021). NS neurons in dlPFC and ACC correspond to putative interneurons and are well separated from BS neurons (**Figure 3A**). In the striatum NS neurons are mostly putative fast spiking interneurons and are well separated from striatal BS neurons (**Figure 3B**) which encompass mostly medium spiny interneurons. Overall, we found only marginal changes in firing rates with Guanfacine that were limited to BS neurons and that were non-systematically distributed (**Figure S2A**). Firing rates were reduced for BS neurons in dlPFC in the attention epoch (p = .034) and in striatum in the feedback epoch (p = .008). Guanfacine did not alter firing variability (coefficient of variation, CV), or the regularity of local interspike intervals (measured as the local variability, LV) in the attention or feedback epoch (**Figure S2B**). We also tested whether Guanfacine affected correlations of spike counts between neurons and found no changes in dlPFC or striatum, but moderately reduced correlations in ACC during the feedback (p < .001) and attention epoch (p < .001) (**Figure S2C**). The reduced firing correlations are consistent with enhanced phasic noradrenergic activation (**Supplemental Discussion**).

### Guanfacine enhances neural encoding of reward prediction errors and trial outcomes

Reversing attention sets relies on recognizing when previously rewarded choices lead to unexpected errors indicating a reversal. We tested this first by quantifying how Guanfacine modulated the encoding of errorneous versus correct outcomes. The correlations of firing rates with outcome rose similarly fast but stronger in the Guanfacine than in the vehicle session in dlPFC, ACC and in the striatum (dlPFC: overall: p = .005 and for post-learning trials p = .002; ACC: for post-learning trials, p = .030; striatum: overall: p = .010; for positive correlations only: p = .014) (**Figure 4B**, see also **Figure S3**). The enhanced outcome encoding reflects an on average increased firing to erroneous outcomes in the dlPFC, ACC, and striatum (average firing to error for Guanfacine and vehicle: 2.8 (±0.5) Hz / 2.7 (±-0.2) Hz, 3.3 (±0.6) Hz / 3.9 (±0.3) Hz, and 3.4 (±1.0) Hz / 3.1 (±0.4) Hz, and to correct outcomes: 2.6 (±0.5) Hz / 2.4 (±0.2) Hz, 3.3 (±0.7) Hz / 3.6 (±0.3) Hz, 3.1 (±0.7) Hz / 2.9 (±0.4) Hz for ACC, dlPFC, and striatum. respectively) (**Figure 2A**). We quantified the difference in [firing rate X outcome] correlations for Guanfacine minus vehicle for outcomes of trials during and after learning completed and for the current and previous trials. We found that Guanfacine increased the correlations with the error versus correct outcomes of the current trial in each of the recorded brain areas during learning, i.e. in trials prior to reaching the learning criterion (dlPFC, ACC and striatum, each: p<0.05) (**Figure 2B**). This finding suggests that Guanfacine may not only encode outcomes, but that it directly modulates the strength of signaling how unexpected an error is, which is quantified as reward prediction errors (RPEs) which serve as learning signals in reinforcement learning models (Hassani et al., 2017). Neurons in dlPFC, ACC, and the striatum encode RPEs during learning and neurons with stronger RPE signals in these areas predict stronger encoding of the reversed target feature after learning (Oemisch et al., 2019). Guanfacine may thus enhance attentional set shifting by modulating RPE encoding. We tested this suggestion and found that Guanfacine enhanced how strong firing rates in the feedback epoch of the task correlated with negative RPEs in the ACC (overall: r = 0.05, p = .039; restricted to the trials prior to reaching learning criterion: r = 0.11, p = .029; negative correlation r = 0.13: p = .016), with positive RPEs in the striatum (overall: r = 0.14, p = .049; with positive RPEs when limited to prior to reaching learning criterion: r = 0.09, p = .016), and for all trials with signed RPEs in the striatum (overall: r = 0.14, p = .024; restricted to the trials prior to reaching learning criterion: r = 0.09, p = .014; negative correlation: r = 0.18, p < .001) (**Figure 2C**).

**Figure 2.**
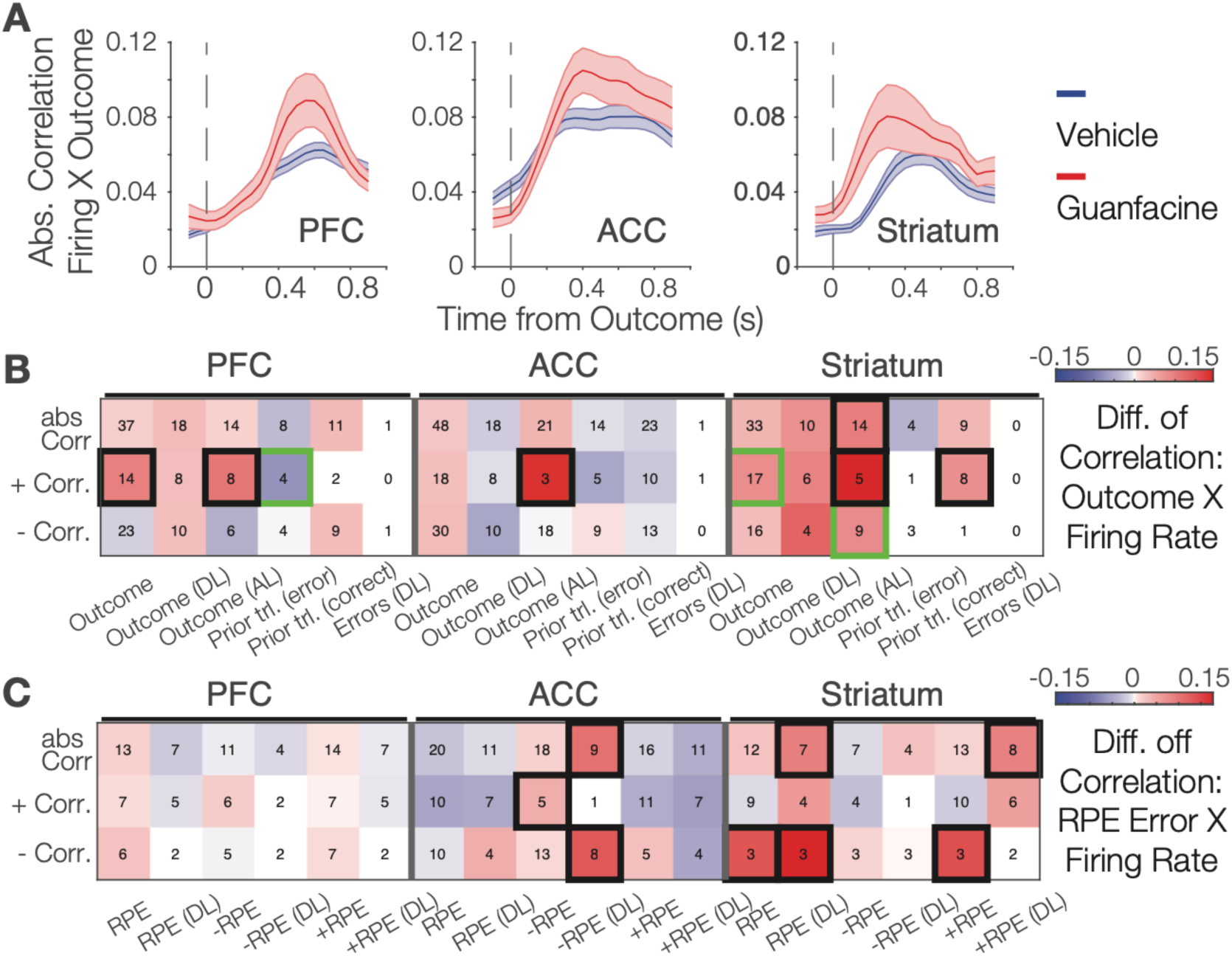
Guanfacine increases correlations of firing rate with reward prediction errors in anterior cingulate cortex and striatum. (**A**) Firing rate correlations with trial outcome (correct vs. error) increase with Guanfacine and vehicle. (**B**) Differences in correlations (Guanfacine minus vehicle) of firing rate in the feedback epoch for different outcome variables (*y-axis*) for neurons record in the dlPFC (left 6 column), the ACC (middle columns) and the striatum (rightmost columns). Stronger colors mean higher (more positive or more negative) correlation coefficients with Guanfacine than vehicle Black squares denote p < 0.05 significance, green squares signify a trend at p < 0.075. Numbers within cells denote the number of neurons in the Guanfacine condition. (**C**) Same format as *A* for firing correlations with reward prediction error variables.

The effects of Guanfacine on outcome encoding and RPE signaling were primarily a modulation of the strength of encoding without altering the overall proportions of neurons encoding outcome variables in the feedback epoch (**Figure S4**).

### Guanfacine enhances outcome encoding particularly for putative interneurons

Previous studies have shown that reversal learning recruits proportionally more narrow spiking (NS) neurons in ACC, dlPFC and striatum (Oemisch et al., 2019; Boroujeni et al., 2020, 2021). We tested whether Guanfacine’s effects on neural coding varied by functional cell type and found that NS neurons (**Figure 3C**), but not BS neurons (**Figure 3D**), encoded outcomes stronger with Guanfacine during the feedback epoch in each of the three recorded brain area. With Guanfacine NS neurons encoded outcomes stronger in dlPFC (unsigned correlations: p = .016; positive correlations: p = .004), in ACC (negative correlations: p = .027), and in the striatum (unsigned correlations: p = .001; positive correlations: p < .001; negative correlations: p = .006). The significant outcome encoding was also evident when the analysis was restricted to trials after learning criterion was reached with stronger outcome encoding after learning in dlPFC (p < .001), ACC (p = .027), and striatum (p = < .001; for negative correlations: p < .001). In contrast to NS neurons, coding of outcomes of BS neurons were unchanged in ACC and striatum and showed reduced encoding of trial outcomes in the dlPFC (p = .016) (**Figure 3D**). There were no neural coding differences of putative neuron types for stimulus or model variables (data not shown).

**Figure 3.**
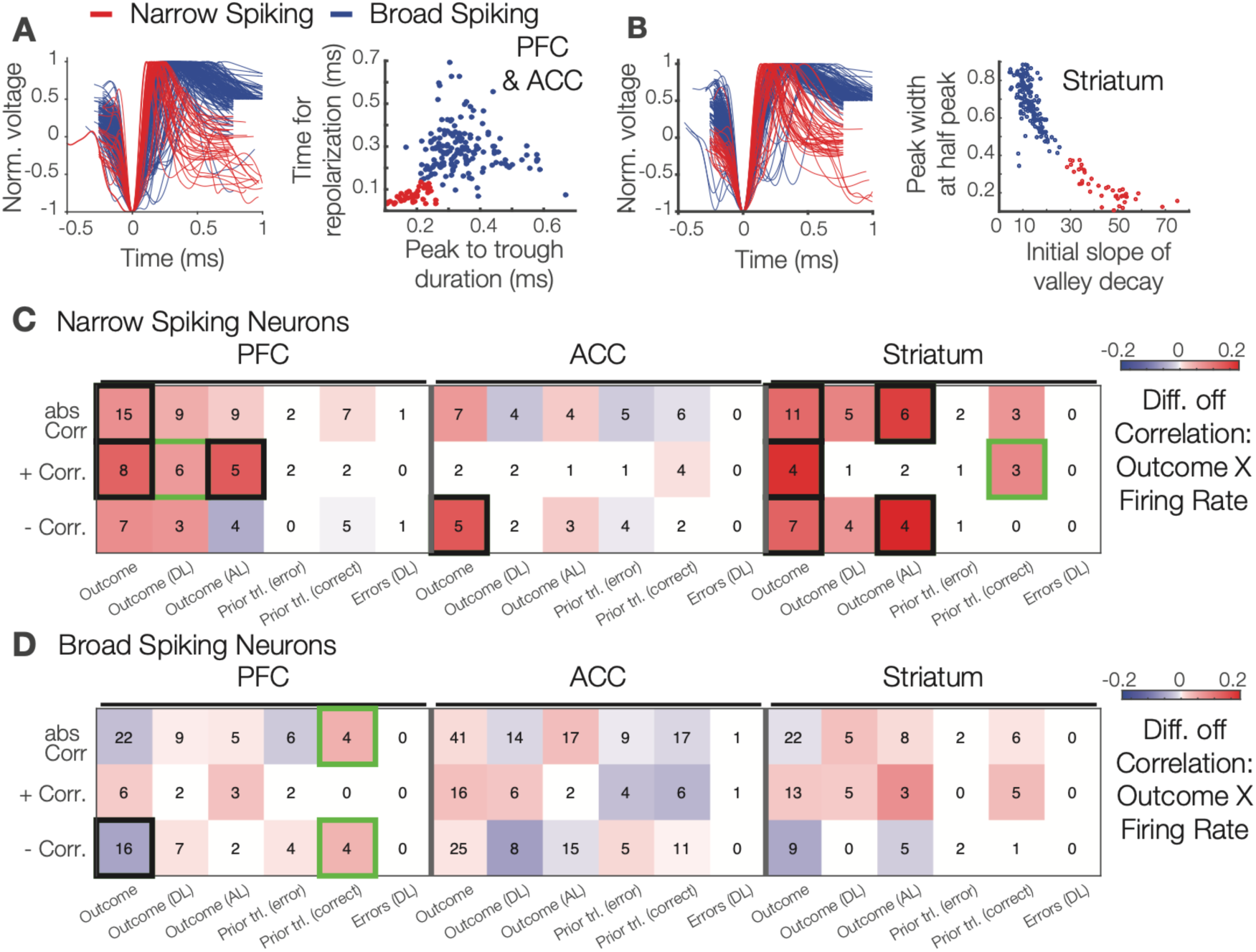
Narrow spiking neurons show stronger firing correlations with outcome variables than broad spiking neurons in dlPFC, ACC and striatum. (**A**) Normalized action potential shapes (*left*) of narrow (red) and broad (blue) spiking neurons recorded in dlPFC and ACC. *Right:* Time to repolarization and peak-to-trough durations distinguishes narrow from broad spiking neurons. (**B**) Normalized action potentials for narrow and broad spiking neurons recorded in the striatum (*left*) distinguished by their spikes’ peak width at half peak and initial slope of valley decay (*right*). (**C**) Differences in correlations (Guanfacine minus vehicle) of firing rate and different outcome variables (*y-axis*) for neurons record in the dlPFC (*left 6 column*), the ACC (*middle columns*) and the striatum (*rightmost columns*).

### Guanfacine enhances the strength and not the prevalence of coding during the feedback epoch

In addition to modulating RPEs and outcome encoding, Guanfacine also increased the encoding of the chosen stimulus color in the dlPFC (p = .003), the chosen location in the striatum (p = .019) and the target location in the ACC (p = .024) in the feedback epoch of the task (**Figure 4**). These effects were evident in example neurons, e.g. for the encoding of the target color and the chosen color (**Figure 4A**), and statistically reliable at the population level (**Figure 4B**). Guanfacine also reduced encoding of the motion direction of the chosen stimulus in dlPFC (p = .032). These changes in Guanfacine compared to vehicle sessions affected primarily the strength of encoding without apparent changes of the relative ranking of which task or model variables were encoded in dlPFC, ACC, and striatum. This conclusion is based on identifying for each neuron the best explaining variable through the regression R^2^ values and ranking neurons according to the best explaining variables, which showed that in the feedback epoch the best-fit variable rankings for Guanfacine and vehicle were significantly correlated and thus similar during the feedback epoch in the dlPFC (p = .006; Tau = .477), ACC (p = .014; Tau = .425), and striatum (p = .014; Tau = .425) (**Figure S5A**).

**Figure 4.**
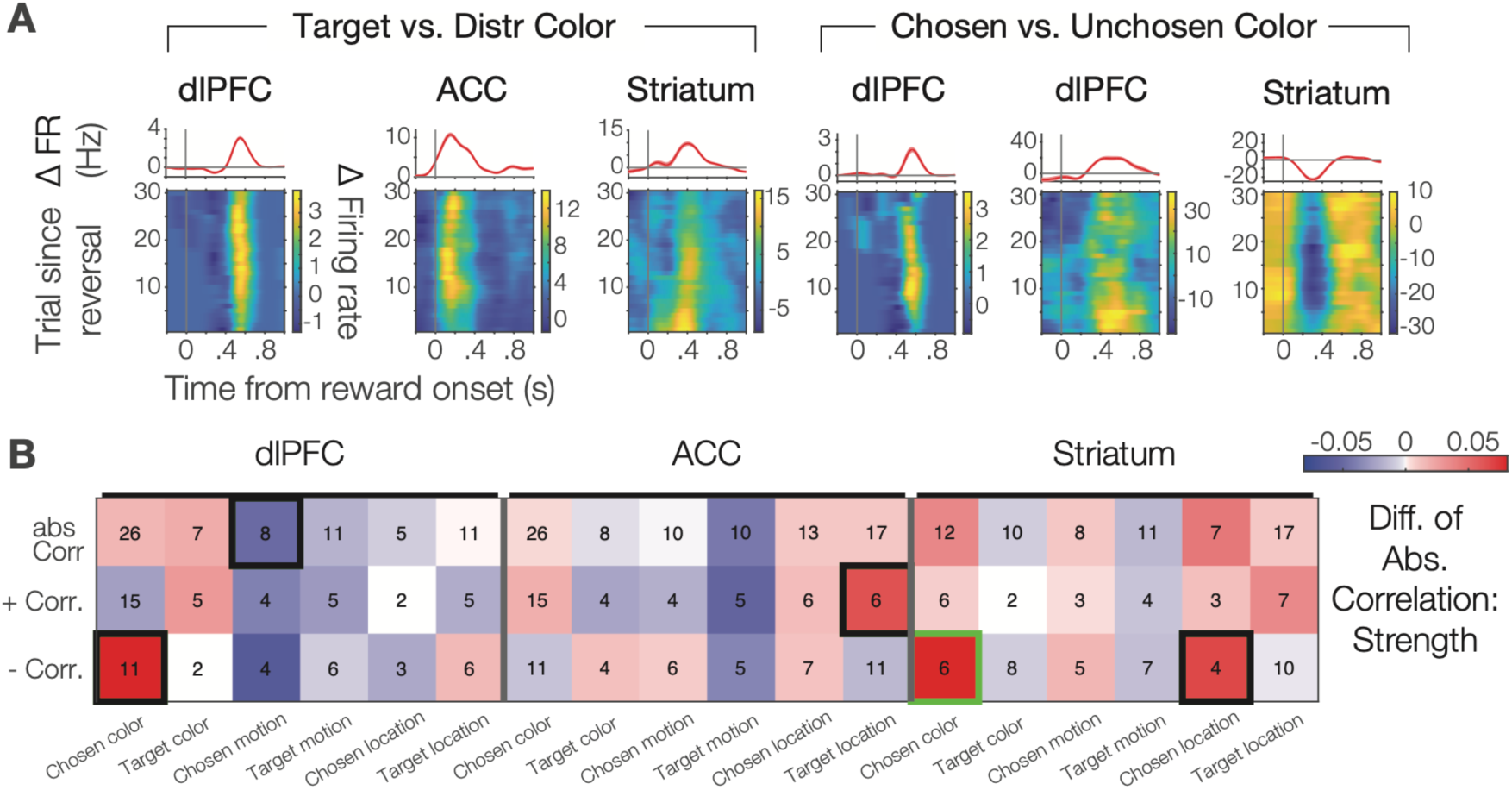
Guanfacine affects the encoding of the target stimulus, and the encoding of color, motion direction and location of the chosen stimulus in the feedback epoch. (**A**) Example neurons showing the firing rate differences for the target versus distractor stimulus (leftmost 3 columns) and for the chosen versus unchosen stimulus color (rightmost three columns). Top panels show firing rate averaged across all trials, while bottom panel colormaps who firing rates across trials (y-axis) and time to reward (x-axis). (**B**) Differences in correlations (Guanfacine minus vehicle) of firing rate and six variables about the chosen stimulus (color, location, motion direction) and target features (color) (*y-axis*) for neurons record in the dlPFC (*left 6 column*), the ACC (*middle columns*) and the striatum (*rightmost columns*). Thick outlined squares denote significant (p<0.05) difference.

### Guanfacine enhances neural encoding of task-relevant color during the attention epoch

A prior study suggested that Guanfacine can enhance in the dlPFC the neural representations of attended, short-term memorized stimulus locations during a delay period prior to making a choice (Wang et al., 2007). In our feature-based set shifting task prior to making a choice, subjects covertly shifted attention to one of two stimuli and sustained attention to the color and location of a stimulus until the Go-signal (a transient stimulus dimming) occurred. During this attention epoch we found that Guanfacine significantly enhanced neuronal encoding of the target color in the ACC specifically for NS neurons (overall: p = .006; for positive correlations: p < .001; for negative correlations: p = .021; **Figure S6A**), but not for BS neurons (n.s.; **Figure S6B**). Consistent with this effect, Guanfacine altered in the attention epoch the ranking of the best-coded variables as reflected by the absence of a correlation of the best-fit rankings of coded task variables between Guanfacine and vehicle conditions in ACC (n.s., Tau = -.281) dlPFC (n.s., Tau = -.007), and striatum (n.s., Tau = .242) (**Figure S5B**). With Guanfacine, in ACC, neurons preferentially encoded the color of the target stimulus, while dlPFC neurons encoded the target color as their second most preferred variable. Outcome variables were encoded non-systematically differently during the attention epoch (**Supplemental Results**).

## Discussion

Here, we reported the behavioral and neural activity signatures of noraderenrgic α2A receptor-specific enhancement of flexible learning. We developed a feature-based attentional set shifting task that required reversing the target color of an attention set through trial-and-error learning. A previous behavioral study suggested that Guanfacine facilitates reversing attentional sets through two mechanisms, enhancing how strong prediction errors are used to update expected values and by reducing the influence of non-attended features (Hassani et al., 2017). Here, we first successfully replicated in two subjects the behavioral effect by showing Guanfacine increased the speed of reversing attention sets and enhanced post-error improvements of performance (**Figure 1D,E**). Secondly, we found neuronal correlates of faster set shifting in the ACC, where Guanfacine enhanced negative RPE signaling, and in the striatum, where Guanfacine enhanced positive RPE signaling (**Figure 3**). Enhanced encoding of RPEs was accompanied by increased firing rate differences of erroneous over correct outcomes in ACC, striatum, and the dlPFC (**Figure 2, 3**).

These finding document a neuronal modulation of outcome and prediction error signaling that lead to faster set shifting by facilicating the updating of expected values after a reversal. Consistent with this scenario we found that Guanfacine also increased neuronal encoding of the target feature color in both, dlPFC and in ACC (**Figure 4**) specifically during the attention period of the task when attention is covertly focused onto the target stimulus.

Taken together, these findings suggest that noradrenergic signaling in all three recorded brain areas, ACC, dlPFC, and striatum, supports cognitive flexibility by enhancing prediction error signaling and improving the representation of the target feature in a top-down attention set. This conclusion is consistent with recent causal manipulation studies in rodent (McBurney-Lin et al., 2022; Su and Cohen, 2022), and calls upon a refinement of frameworks that propose a more general involvement of noradrenergic signaling to either enhance working memory for task relevant stimulus representations (Wang et al., 2007), or track the uncertainty (including the unexpecteness of outcomes) of an environment and facilitate exploratory behaviors by reducing uncertainty (Yu and Dayan, 2005; Dayan and Yu, 2006).

### Guanfacine enhances prediction error signaling in ACC and striatum

Our findings of enhanced RPE encoding and faster reversal learning support conclusions from studies in mice documenting noradrenergic neurons in the locus coeruleus causally support flexible behavioral switching of stimulus – reward associations (McBurney-Lin et al., 2022; Su and Cohen, 2022). In these studies, activity of locus coeruleus neurons were either necessary for faster learning (Su and Cohen, 2022) or were predictive of faster behavioral switches (McBurney-Lin et al., 2022). The results reported here suggests that these noradrenergic signals in the LC are translated into stronger prediction error encoding along the ACC-striatum pathway. According to this suggestion noradrenergic inputs affect those ACC and striatum neurons that encode reward prediction errors and can be boosted either by activating the locus coeruleus (in the rodent studies), or by increasing α2A noradrenergic activation pharmacologically with Guanfacine to facilitate synaptic activity of neurons encoding RPEs in the ACC and striatum. Our results predict that this modulation is specific to the subset of neurons in ACC and striatum that encode outcomes and RPEs, because Guanfacine did neither affect the overall firing rates of neurons (**Figure S2A**,**B**), nor did it change the prevalence of neurons encoding other task variables during the processing of outcomes (**Figure S5**). According to this interpretation Guanfacine modulates the intrinsic neuronal signaling of each these brain areas without recruiting additional neuronal populations.

### Guanfacine enhances signal-to-noise ratio of neural signaling

Guanfacine enhanced the correlation of neural firing with task relevant variables without increasing the number of neurons with firing correlations. This pattern of results reflects an enhanced neuronal signal-to-noise ratio for coding task relevant variables, which included better encoding of the color feature of the target stimulus. This conclusion corroborates and extends the seminal NHP study of Guanfacine in dlPFC by Arnsten, Wang and colleagues (Wang et al., 2007). In their study locally injected Guanfacine enhanced persistent delay period neural firing in dlPFC for those spatial locations that were preferred by the recorded neurons. Enhanced firing for preferred over nonpreferred spatial locations indexes an enhanced signal-to-noise ratio of short-term memory representations. Our study extends these insights from the dlPFC supporting delayed-match-to-sample performance to the ACC and striatum supporting feature-based attentional set shifting performance. The converging conclusions in both studies suggest that α2A receptor activation has similar effects of enhancing the gain of neuronal responses in different brain systems. Gain modulation has been linked to noradrenergic activation, capable of potentiating responses of activated neurons that are already recruited by ongoing task demands (Berridge and Waterhouse, 2003; Aston-Jones and Cohen, 2005; Ferguson and Cardin, 2020). Taken together the evidence from Wang et al., and our study predicts that α2A receptors may enhance information processing beyond the ACC, dlPFC and striatum also for other brain areas and for task variables that are preferentially encoded in a specific area. In the dlPFC the enhanced signa-to-noise ratio firing could be traced back to the activation of post-synaptic α2A receptors on dendritic spines of basal dendrites of pyramidal cells in the dlPFC leading to sustained firing through a reduced intracellular cAMP signaling (Arnsten et al., 2010; Cools and Arnsten, 2022). Whether this mechanism of postsynaptic α2A receptors on dendritic spines of pyramidal cells will be specific to the dlPFC and working memory processes, or whether it can be extended to the ACC and flexible learning, or also apply to noradrenergic modulation in posterior parietal or visual cortices is an important question for future research.

### Guanfacine’s effect on narrow spiking outcome signaling

Our electrophysiological recordings allowed distinguishing narrow spiking (NS) from broad spiking (BS) neurons (**Figure 3A**), with NS neurons containing fast spiking putative interneurons. We found that Guanfacine enhanced outcome encoding specifically for these putative interneurons in each of the three recorded brain areas but had no effect on BS neurons (**Figure 3B**,**C**). The clarity of this finding was unexpected and is intriguing, suggesting that the α2A agonist may selectively modulate outcome signaling of putative interneurons. This interpretation is consistent with studies documenting that adrenoceptor expression and modulation is stronger for interneurons than pyramidal cells with α2 and β adrenoceptors enhancing their inhibitory actions while α1 adrenoceptors decrease their inhibitory actions in prefrontal cortex (Kawaguchi and Shindou, 1998; Wang et al., 2013; Liu et al., 2014; Xing et al., 2016; Lee et al., 2020), as well as in sensory and sensorimotor cortices (Bennett et al., 1998; Nai et al., 2009; Salgado et al., 2011, 2012; Ohshima et al., 2017). Thus, Guanfacine may enhance the inhibitory tone in neural circuits. Our findings suggest that this effect was not translated into changes in overall firing, because NS neurons had unaltered firing with Guanfacine and BS cells showed only marginal and non-systematic reductions of BS firing. An alternative scenario is that Guanfacine enhanced the neuronal gain of inhibitory interneurons (Ferguson and Cardin, 2020), which could explain stronger outcome encoding of NS neurons not only during set shifting but also after learning a new attention set was completed. However, we cannot be certain whether this mechanism could also underlie enhanced learning as we had not enough NS neurons for analyzing correlations of NS neurons only in trials during learning. Future studies will need to test specifically whether the α2A receptor is critical for the prominent role of putative NS interneurons to predict learning success and encode prediction errors in ACC, dlPFC and striatum (Oemisch et al., 2019; Boroujeni et al., 2020, 2021).

### Alpha 2A receptor specific improvement of flexible set shifting

Our study documents that Guanfacine enhances flexible reversal learning of attention sets, which adds clarity to the potential of Guanfacine as a highly selective α2A acting drug to enhance attentional functions (Berridge and Waterhouse, 2003; Aston-Jones and Cohen, 2005; Berridge and Spencer, 2016). Our results indicate that the attentional benefit of Guanfacine includes a better recognition of erroneously attended stimuli as indicated by enhanced post-error improvement (**Figure 1E**). The post-error adjustment effect was limited to the exploratory learning period immediately after a reversal. During the plateau performance period, Guanfacine did neither modulate post-error adjustment nor overall accuracy levels. Thus, Guanfacine specifically improved the rate of learning to reverse the attention sets, consistent with it creating a new attentional state (Sadacca et al., 2017). This conclusion also resonates with prior modeling studies of Guanfacine (Hassani et al., 2017) and with a number of human psychopharmacological studies that have associated the action of NE/dopamine reuptake inhibitors with the modulation of learning rates based on environmental uncertainty (Jepma et al., 2016; Howlett et al., 2017; Cook et al., 2019).

We found the neural correlates of enhanced set shifting in those neural circuits in the ACC and striatum of NHPs that are known to activate during guided exploratory behavior and information seeking (Bromberg-Martin and Monosov, 2020; Boroujeni et al., 2022; Jahn et al., 2023). Activity in the ACC, in particular, correlates with successfully exploring alternative options and gathering information about the value of options (Jahn et al., 2023). In the feature-based set shifting task used here, guided exploration during the reversal period will help to overcome previous response biases when updating attentional sets (Sadacca et al., 2017).

Previous studies with α2A agonists in NHP have overwhelmingly described working memory improvements (reviewed in Hassani et al., 2017), while rodent studies have included tasks testing reversal learning and attention set shifting and have produced complex results (e.g. Lapiz and Morilak, 2006). While our study documents that α2A stimulation is sufficient to enhance learning flexibility, its means to modulate circuits might also involve interactions with dopamine (Cools et al., 2009; Stelzel et al., 2013; Varazzani et al., 2015) (**Supplemental Discussion**).

### Possible implications beyond reversing attention sets

Previous studies suggest that noradrenergic tone modulates how optimal rats exploit foraging patches. Noradrenergic tone can predict whether mice show exploratory versus exploitative behaviors (Koralek and Costa, 2021) and overactivation of the locus coeruleus increased exploratory behaviors sub-optimally, causing animals to leave reward-depleting patches earlier than is optimal (Kane et al., 2017). The mechanisms for suboptimal exploratory foraging behavior are largely unknown, but our study provides some clues that noradrenergic modulation of foraging will involve the speed with which expected values are updated. With a good tonic noradrenergic activity state, reward prediction errors are efficiently signaled, and object values effectively updated. We expect that at non-optimal tonic concentrations of noradrenaline that these value updating processes are impaired, leading either to slower recognizing when object values change in an environment or to faster updating of values leading to faster shifting of behavior. We therefore predict that the faster leaving of depleting patches with enhancing locus coeruleus tonic activity reflects a more rapid updating of value expectations leading to an earlier disengagement with the current patch (Kane et al., 2017).

In summary, our findings illustrate that Guanfacine can improve adaptive, flexible updating of attention sets by increasing the efficiency of prediction error signaling in medial prefrontal and striatal circuits. These insights suggest that α2A receptor activation across the fronto-striatal network is a key player to mediate noradrenergic regulation of behavioral flexibility.

## Materials and Methods

### Subjects and apparatus

Data was recorded from two male rhesus macaques (*Macaca mulatta*) age 7 and 9 years old. Both subjects were fluid restricted during the length of the experiment with unrestricted access to chow. All animal care and experimental protocols were approved by the York University Council on Animal Care and were in accordance with the Canadian Council on Animal Care guidelines.

Animals were seated in a custom-made primate chair and placed in a dark, sound attenuated booth such that their eyes were 65 cm away from a 21’ LCD monitor with a 85 Hz refresh rate. Experimental control, including stimulus presentation, eye positioning monitoring and reward delivery was done through MonkeyLogic (open-source software https://www.brown.edu/Research/monkeylogic/). Eye positions were calibrated and tracked monocularly using a video-based eye tracking system (Eyelink 1000 Osgoode, Ontario, Canada; 500 Hz sampling). Eye calibration occurred daily using a 9-point fixation pattern and was monitored throughout each session. Liquid reward, controlled from an air-pressure mechanical valve system (Neuronitek, London, Ontario, Canada) was delivered via a sipper tube.

### Behavioral task

The animals performed a feature-based reversal learning task as previously described (Hassani et al., 2017; Oemisch et al., 2019). Subjects learned through trial-and-error which one of two grating stimuli was deterministically rewarded in any given block lasting at least 30 trials or until a 90% performance criterion over the preceding 12 trials was reached) (**Figure 1A**). Each grating stimulus was defined in any given trial by a combination of three features: location (left vs right), color (monkey Ha: red vs green; monkey Ke: cyan vs yellow), and motion direction of the stimulus grating (up vs down). The two stimuli always contained opposite values for each of these three dimensions. Only color was indicative of reward value and thus is referred to as the attention cue in text, while location and motion were randomly associated with reward.

Trials proceeded through a motion and color cue period before a transient dimming of the target stimulus instructed the subjects to make a saccadic choice, while a dimming of distractor stimuli had to be ignored (**Figure 1B**). In particular, a trial started with the appearance of a gray central fixation point, which the subjects had to fixate. After 0.5–0.9 s, two black/white gratings appeared to the left and right of the central fixation point. Following another 0.4 s, the two stimulus gratings gained a color or had their gratings drift in opposite directions (up or down), followed after 0.5–0.9 s by the onset of the second stimulus feature such that both stimuli eventually had both color and motion. After 0.4–1 s, the two stimuli dimmed simultaneously for 0.3 s or one stimulus dimmed first followed by the other separated by 0.55 s. The dimming represented a go-cue to make a saccade from the central fixation point to one of two response targets displayed above and below the central fixation point. Breaking fixation from the central fixation point before the dimming event of the target terminated the trial. In order to acquire reward, subjects had to make either an upward or downward saccade matching the direction of motion of the stimulus grating with the rewarded color, and this saccade had to occur no later than 0.55 s after the dimming of the stimulus with the rewarded color.

### Statistical measure of learning

To identify learned blocks and an individual trial where statistically reliable learning could be said to have occurred in each block, we used an ideal observer estimation maximization (EM) algorithm (Smith and Brown, 2003; Smith et al., 2004). Briefly, this framework utilized a state equation to represent the internal learning process as a hidden Markov or latent process which was updated with each trial. This provided an estimate of the probability of a correct choice taking into account all trials within the block (**Figure 2A** *bottom*). The learning trial was then defined as the trial during which the lower 95% confidence bound exceeded chance (p = 0.5) and did not drop back down below chance for the rest of the block.

### Drug dosing

Guanfacine was purchased (Guanfacine hydrochloride; Sigma-Aldrich, St. Louis, MO) and prepared with sterile water vehicle (0.1 mL volume) immediately before blinded IM injections. Subjects received Guanfacine (0.075 mg/kg) or sterile water vehicle injections close to 2 hours before the start of the first trial (mean: 135 ± SE 2 min). Each week contained at most a single Guanfacine administration day which was always either on Thursday or Friday while vehicle data was collected on either Tuesdays or Wednesdays; animals still trained every day. In total, we recorded 17 and 12 Guanfacine days for monkey Ha and monkey Ke respectively. We the selected dose of Guanfacine because it was previously shown that it enhances performance in this task (Hassani et al., 2017).

### Electrophysiological recordings and unit isolation

Single contact tungsten electrodes (FHC, Bowdoinham, ME; 1.2-2.2 MOhm impedance electrodes) were used for extracellular recordings. They were loaded into up to 4 software-controlled precision micro-drives (NAN Instruments Ltd., Israel) and lowered into the brain through a 20x25 mm rectangular recording chamber guided by MR images. Single units were recorded in the dlPFC (area 46), the ACC (area 24), and the head of the caudate nucleus (Calabrese et al., 2015) (**Figure 1D**). Data amplification, filtering and acquisition were done with a multi– channel acquisition processor (Neuralynx). Spiking activity was obtained following a 300-8000 Hz passband filter and further amplification and digitization at 40 kHz sampling rate. After the initial acquisition of highly isolated waveforms in the regions of interest, electrodes were left to stabilize for 30-60 minutes before the start of the task. Sorting and isolation of single unit activity was performed manually offline with the Plexon Offline Sorter, based on principal component analysis of the spike waveforms. In order to maximize statistical power in neural analyses, an extended dataset previously recorded from monkey Ha without any injections was also considered and pooled with the vehicle data and referred to as ‘non-drug data’. Although behavioral performance in these sessions was superior to the vehicle sessions, virtually all relevant behavioral trends and results remained consistent (data not shown).

### Putative cell type classification

Highly isolated single units were classified based on the properties of their actin potential waveforms using previously published methods (Lansink et al., 2010; Ardid et al., 2015; Oemisch et al., 2019). Briefly, all waveforms were normalized and aligned to their peak and classified by clustering the first PCA of different metrics based on if they were cortical or striatal in origin (**Figure 6C-D**). Cortical neurons (from dlPFC or ACC) were classified based on peak-to-trough duration and time to repolarization with broad spiking neurons being putative pyramidal neurons and narrow spiking neurons being putative interneurons. Striatal neurons (from striatum) were classified based on peak width and the initial slope of the valley decay with broad spiking neurons being putative medium spiny neurons (MSNs) and narrow spiking neurons being putative fast spiking interneurons (FSIs).

### Multi-linear regression

Spike trains were transformed into spike-density functions smoothed with a Gaussian kernel with a standard deviation of 50 ms. Only correct (rewarded) and incorrect choice trials were analyzed; incorrect trials were defined as unrewarded trials where either the unrewarded object was chosen or any choice was made during the dimming (go signal) of the unrewarded object (either before or after the dimming of the rewarded object). The average trial-wise activity during the epoch of interest (0.05-1s during the feedback epoch and 0.05-0.7s during the attention cue onset epoch) of each neuron was regressed to 18 variables that were classified as either stimulus variables, outcome variables or latent model variables. The 6 binary stimulus variables were the color (color 1 vs color 2), motion (up vs down) and location (left vs right) of the chosen stimulus and the color, motion and location of the rewarded stimulus. The outcome variables were trial outcomes (binary: rewarded or unrewarded), trial outcomes during learning only (see **Statistical measure of learning** above), trial outcomes after learning, prior trial outcome for correct trials (binary: rewarded trial preceded by a rewarded trial or an error trial), prior trial outcome for error trials (binary: error trial preceded by a rewarded trial or an error trial) and error trial order during learning (non-binary: ranking errors in descending order until the statistically defined learning trial). The latent model variables were all non-binary but different depending on the epoch in question. During the feedback epoch, the latent model variables were signed RPEs, positive RPEs, negative RPEs, and the same three variables for trials during learning only. During the attention cue onset epoch, the latent model variables were the choice probability of the chosen stimulus, value of the chosen rewarded stimulus, value of the chosen unrewarded stimulus and the same three variables but for trials during learning only.

A single neuron may have multiple significant regressions and for each significant regression (neurons had to be isolated for at least 30 trials), a correlation coefficient was computed. These coefficients were then averaged for Guanfacine and non-drug neurons and their difference plotted separated between unsigned, positive and negative correlation coefficients for each brain region and also split by putative cell types. Statistical comparison between Guanfacine and non-drug neurons was done through bootstrapping with shuffled condition labels (5000 permutations). Only comparisons with at least 3 neurons in both Guanfacine and non-drug categories were statistically tested.

For each neuron, the strongest regression (significant regression with the highest R^2^) was also identified. Then the variables that best explained activity in each brain region were ranked based on the proportion of neurons that had the highest R^2^ value per variable (Padoa-Schioppa and Assad, 2006). This ranking was done separately for Guanfacine and non-drug days which were then compared using Kendall’s tau correlation.

### Model variables

We estimated latent variables underlying learning performance including the reward prediction error and the expected stimulus values using a hybrid Bayesian-reinforcement learning model as validated and described in previous studies (Niv et al., 2015; Hassani et al., 2017; Oemisch et al., 2019; Womelsdorf et al., 2021). This model was the best performing model among a number of different reinforcement learning, Bayesian and hybrid models (see Hassani et al., 2017 for detailed model description and comparisons). Briefly, the model describes each object’s value as a weighted combination of its features (color, location, motion). For each trial, a single object is then selected through a softmax selection process with a RPE being computed based on the outcome. This RPE signal is then scaled by a learning rate to adjust the values of the chosen object’s features while the feature values of the unchosen object decay.

## Data availability

The data that are part of this study are available from the corresponding authors upon reasonable request.

## Code availability

The custom-made MATLAB scripts are available from the corresponding author upon reasonable request.

## Acknowledgements

The authors would like to thank Mariann Oemisch and Marzyeh Azimi for their help with animal training and data collection, Paul Tiesinga for being instrumental in developing the reinforcement learning model, and Hongying Wang for help with drug administration and animal care. This research was supported by grants from the Canadian Institutes of Health Research (CIHR) and the National Institute of Mental Health of the National Institutes of Health under Award Number R01MH129641 (TW). The content is solely the responsibility of the authors and does not necessarily represent the official views of the National Institutes of Health.

## Author contributions

SAH conducted the experiment and analysis and contributed to writing the paper. TW conducted analysis, wrote the paper, and supervised the project.

## Materials and correspondence

Requests for correspondence and materials can be addressed to Dr. T Womelsdorf.

