## Supplemental Information for "Noradrenergic alpha-2a Receptor Stimulation Enhances Prediction Error Signaling in Anterior Cingulate Cortex and Striatum"

**Content:**

- **Supplemental Methods**
- **Supplemental Results**
- **Supplemental Discussion**
- **Supplemental Figures S1-S7**

### Supplemental Methods

Pupil diameter

Monocular measurements of pupil diameter were made during experiments by the eye tracker (Eyelink 1000; 500 Hz). The pupil diameter traces were only considered during the 400 ms after stimulus onset and before the onset of either the motion or color features of the object as it was the longest period with the most stable visual stimulation. Only the traces for completed trials in the first three blocks of each session was considered and averaged across the entire 400 ms time bin. These trial averages were then z-normalized by subtracting the grand-averaged pupil diameter of first block trials for each monkey. We then averaged the z-normalized pupil diameter for the first block of sessions with and without Guanfacine.

Spiking activity

Spike trains were transformed into spike-density functions smoothed with a Gaussian kernel with a standard deviation of 50 ms. Only correct (rewarded) and incorrect choice trials were analyzed;

incorrect trials were defined as trials where the unrewarded object was chosen or any choice being made during the dimming (go signal) of the unrewarded object (either before or after the dimming of the rewarded object). Then the firing rate during the epoch of interest (feedback epoch: 50-1000 ms after feedback; attention cue onset epoch: 50-700 ms after color onset) across all trials was averaged for each neuron and then neurons with or without the administration of Guanfacine were then compared with permutation testing where the condition (Guanfacine or not) was randomly shuffled.

The regularity of spikes was calculated for each neuron using the measures of coefficient of variation (CV) and local variability (LV) (Shinomoto et al., 2003, 2005). Both CV (formula 1) LV (formula 2) were computed over the entire length of all completed trials using inter-spike intervals (T) where $\bar{T}$ is the mean inter-spike interval.

$$CV = \sqrt{\frac{1}{n-1}\sum_{i=1}^{n}(T_i - \bar{T})^2} \Big/ \bar{T} \quad (1)$$

$$LV = \frac{1}{n-1}\sum_{i=1}^{n-1}\frac{3(T_i - T_{i+1})^2}{(T_i + T_{i+1})^2} \quad (2)$$

The spike count pair-wise correlation (Rsc) was computed by adapting previously described methods (Joshi and Gold, 2022). Spike counts were calculated over the entire epoch of interest (feedback epoch: 50-1000 ms after feedback; attention cue onset epoch: 50-700 ms after color onset) on every trial. For each neuron pair from the same brain region (dlPFC: dorsolateral prefrontal cortex, ACC: anterior cingulate cortex, and striatum), only trials where both neurons were stably isolated were considered. The spike count for each neuron was z-scored with trials that were > 3 standard deviations outside of the mean being excluded from both neurons. A Pearson correlation coefficient was then calculated for each neuron pair and separated based on if it were not significant, significant and positive or significant and negative. The correlation coefficients (Rsc) were then split between Guanfacine and non-Guanfacine days where their median, as well as a 95% confidence interval was extracted through bootstrapping and plotted (**Figure S2C**). Guanfacine and non-Guanfacine Rsc was then compared in the feedback epoch and the choice feature onset epoch by a Wilcoxon rank sum test with Bonferroni correction.

### Supplemental Results

#### Pupil diameter

Although pupil diameter is often used to infer details about locus coeruleus (LC) activity, it is also highly related to the activity in many other brain regions as well (Murphy et al., 2014; Joshi et al., 2016; Reimer et al., 2016; Joshi and Gold, 2022). Even when only considering the LC however, it can be highly variable especially between sessions (Megemont et al., 2022). Nevertheless, to see if the systemically administered dose of Guanfacine (0.075 mg/kg) did reduce LC activity, we looked at changes in pupil diameter during the first three blocks of each session where its concentration would be the highest. Pupil diameter was averaged across the 400 ms window after the onset of the two graded stimuli and before the onset of either the color or motion features. Due to each session's data having either Guanfacine administration or not, no direct same-session comparisons could be made. Therefore, pupil diameter trial averages for the first three blocks, where Guanfacine concentration is highest, were z-normalized by subtracting the grand-average for each monkey during those trials before comparing Guanfacine administration sessions with vehicle. We found a significant reduction in the first three blocks' pupil diameter with Guanfacine administration for both monkey Ha (t-test; $p < .001$) and monkey Ke (t-test; $p = .010$) (**Figure S1C**).

**Behavioral performance**

Monkey Ha and Ke learned 52.1% and 62.3% of blocks respectively with an average learning trial speed of 15.7 and 18.7 respectively. With Guanfacine, monkey Ha and Ke learned -2.5% and +5.3% more blocks with an average learning speed change of 2.9 and 3.0 trials respectively. The summed difference in reward probability after reversals were significantly different between vehicle and Guanfacine administrations (Ha: $p = .042$; Ke: $p = .021$; **Figure S1**) as were the summed difference in the EM derived reward probabilities (Ha: $p = .035$; Ke: $p = .004$; **Figure S1**).

**Changes in firing properties**

We investigated the impact of Guanfacine on the firing properties of neurons in the dlPFC, ACC and striatum. We looked at both the feedback epoch (50-1000 ms after feedback) and the attention cue onset epoch (50-700 ms after color onset which is the minimum length of this epoch). Guanfacine did not significantly change the firing rate in the dlPFC, ACC or striatum during the feedback epoch (permutation testing: 5000 permutations; all n.s.) or the attention cue onset epoch (permutation testing: 5000 permutations; all n.s.; **Figure S2A**). Separating neurons by their putative cell type revealed a significant reduction in firing rate in broad spiking neurons in the dlPFC and striatum during the feedback epoch and no changes in the attention cue onset epoch (permutation testing: 5000 permutations; dlPFC: $p = .034$; ACC: n.s.; striatum: $p = .008$; not shown).

We also looked at changes in spiking regularity across the whole trial via the CV and LV in the dlPFC, ACC and striatum. Guanfacine administration did not significantly change the CV of neurons in the dlPFC, ACC or striatum although a significant main effect of area was found (Anova: $F(2,1150) = 13.85$; $p = < .001$) with the caudate having a significantly lower CV than the dlPFC (Tukey's: $p < .001$) and the ACC (Tukey's: $p = .004$) (**Figure S2B** *top*). Results did not change when only considering narrow or broad spiking neurons (data not shown). Similarly, no significant change in LV was observed with Guanfacine administration with no significant main effects or interactions (anova: all n.s.). This remained consistent even when only considering narrow or broad spiking neurons (data not shown).

As expected, the correlation of binned firing rate (200 ms centered windows with 50 ms shifts) with outcome during the feedback epoch rose at the time of feedback presentation, reaching some peak within 1000 ms of feedback onset (**Figure S3**)..

### Supplemental Discussion

#### How α2A stimulation relates to locus coeruleus activity

Pre-synaptic α2A adrenoceptors act as noradrenergic auto-receptors, reducing the further release of NE. It has been previously shown that systemic Guanfacine administration reduces LC activity (Engberg and Eriksson, 1991; Okada et al., 2018). Consistent with reduced LC firing we observed reduced pupil diameter in the blocks temporally closest to the inject time (**Figure S1D**). However, Guanfacine also resulted in on average reduced pair-wise spike count correlations in the ACC relative to the non-drug condition (**Figure S2C**), which a recent study has shown to be indicative of higher LC activity (Joshi and Gold, 2022). This discrepancy might be resolved by distinguishing tonic from phasic LC activity modulations. The increases of pairwise firing correlations in Joshi and Gold (2022) likely reflect reduced tonic LC firing, while the reduction of firing correlation that we found indicates enhanced phasic LC firing in the presence of reduced tonic LC firing. This proposal is consistent with findings showing that noradrenergic auto-receptor activation can increase LC neuron sensitivity to glutamatergic and cholinergic stimulation thus emulating an increase in phasic LC firing (Aston-Jones et al., 1991; Aston-Jones and Cohen, 2005b). This suggests that systemic Guanfacine administration may reduce tonic LC activity while simultaneously boosting phasic LC activity.

#### Possible interactions of α2A stimulation with other neuromodulators

Our data suggests that the stimulation of the α2A adrenoceptor is sufficient for enhancing RPE encoding in the fronto-striatal network and more flexible learning. It is important to note, however, that due to the nature of systemic administration we cannot be certain that the observed effects are directly mediated by α2A adrenoceptors because other neuromodulatory systems interact heavily with NE, including strong interactions between dopamine and NE, particularly in the PFC (Devoto et al., 2005; Jentsch et al., 2008; Xing et al., 2016; Cools and Arnsten, 2022), and interactions with 5-HT and acetylcholine (Aston-Jones et al., 1991; Berridge and Foote, 1991; Berridge and Wifler, 2000). Such interactions suggest some involvement of dopamine in affecting cognitive flexibility (Marshall et al., 2016; Xing et al., 2016; Wang et al., 2018), with evidence that prefrontal dopamine is in part provided by LC noradrenergic terminals (Devoto et al., 2005, 2019, 2020). It will be an important venue for future research to understand which other neuromodulatory systems might support cognitive flexibility beyond the α2A adrenoceptors.

### Figure legends

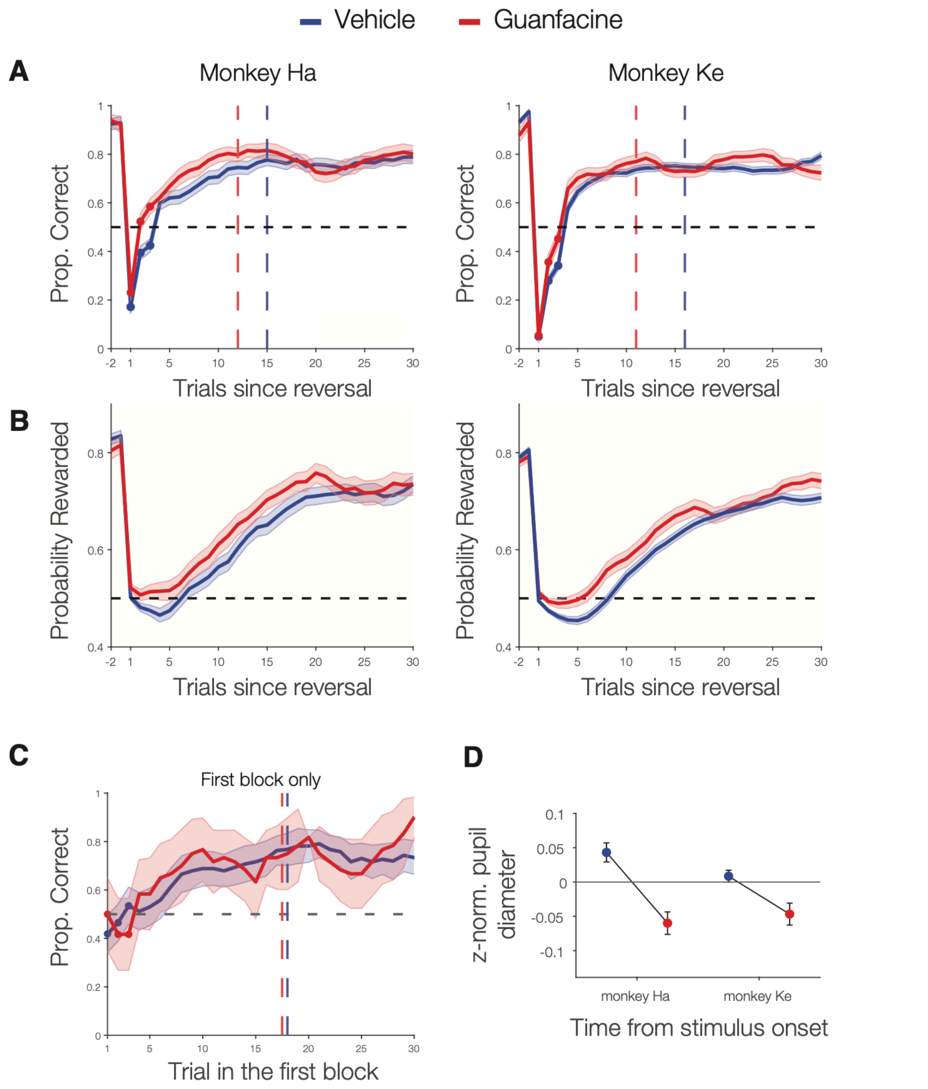


**Figure S1**. Individual monkey performance curves and pupil diameter. (**A**) Increasing proportions of correct choices over trials shows faster learning with Guanfacine than vehicle in both monkeys. (**B**) Ideal observer estimate of the probability to observe rewarded choices for monkeys Ha (*left*) and Ke (*right*). Learning curves were smoothed except for the first 3 trials. The horizontal dashed line represents chance probability. The vertical dashed lines represent the median learning trial for Guanfacine (*red*) and vehicle (*blue*). (**C**) Learning curve of only the first block of each session for Guanfacine (*red*) and vehicle (*blue*). Reversals started by definition only with the second block in each session. (**D**) Average normalized pupil diameter in the first 3 blocks of Guanfacine (*red*) and vehicle (*blue*) sessions.

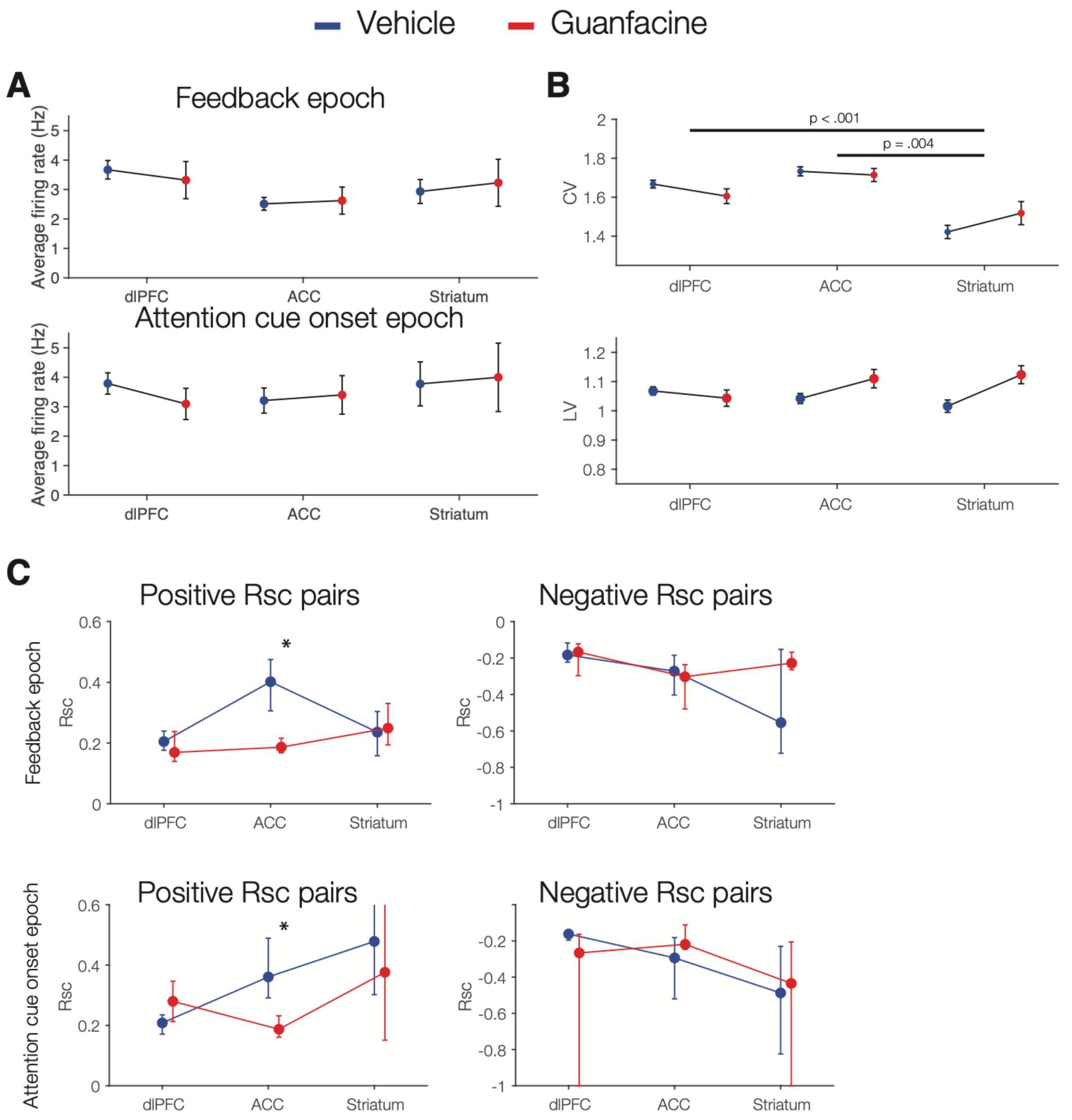


**Figure S2**. Guanfacine had marginal effects on spike timing and pairwise firing correlations. (**A**) Average firing rate during 50-1000 ms after feedback (*top*) and during the covert attention period (50-700 ms after attention cue onset; *bottom*). Firing rate was averaged during the epoch of interest across all completed trials for each neuron and then averaged between all neurons for each brain region and condition. (**B**) The coefficient of variation (CV; *top*) and local variability (LV; *bottom*) of recorded neurons. The LV indexes the regularity of interspike intervals (*see* Methods). Both CV and LV were calculated using all completed trials over the entire trial length. (**C**) The pair-wise spike count correlation (Rsc) during the feedback epoch (*top*) and attention epoch (*bottom*) split between positive Rsc pairs (*left*) and negative Rsc pairs (*right*). Values and error bars represent boot strapped averages and 95% confidence intervals. A Wilcoxon rank sum test with Bonferroni correction was used to compare the Rsc of neuron pairs in Guanfacine and vehicle session.

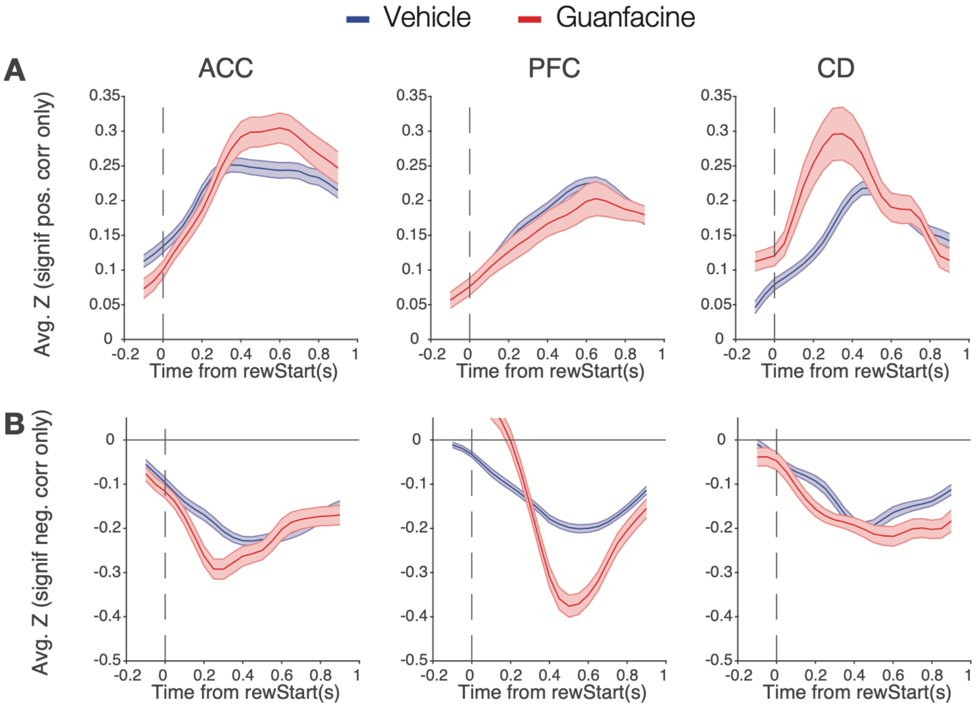


**Figure S3**. Firing rates correlate with outcomes (correct vs error) during the outcome epoch. (**A**,**B**) The firing rate of each neuron was binned in 200 ms windows (50 ms shifts) and correlated with the trial outcome for neurons with a significant positive (*A*) and negative (*B*) correlation for the dlPFC (*left*), ACC (*middle*) and striatum (*right*). Red and Blue denote Guanfacine and vehicle conditions.

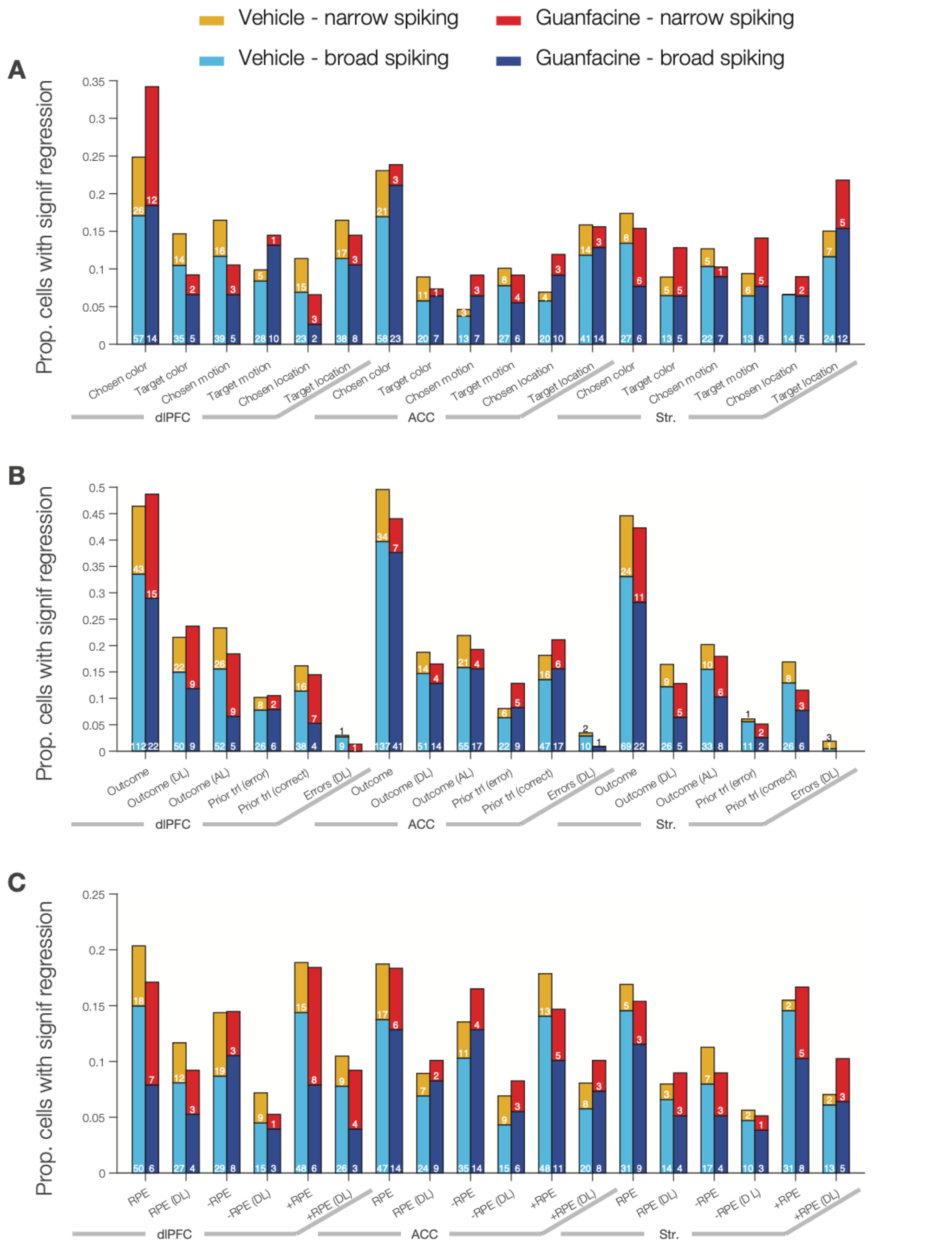


**Figure S4.** The proportion of neurons with significant ($p<0.05$) regression coefficient for firing rates explained by task-relevant variables during the feedback epoch with Guanfacine (*red* and *dark blue*) and during the vehicle condition (*orange* and *light blue*). (**A**) Regression with variables characterizing the chosen stimulus and the target stimulus (its color, motion direction, location) split into narrow spiking neurons (*orange* and *red*) and broad spiking neurons (*light blue* and *dark blue*). The small white or black numbers within the bars denote the actual cell counts. (**B**) Same format as *A* for regressions with variables characterizing the current or previous trials' outcome. (**C**) Same format as *A* and *B* for regressions with latent model-derived reward prediction error variables.

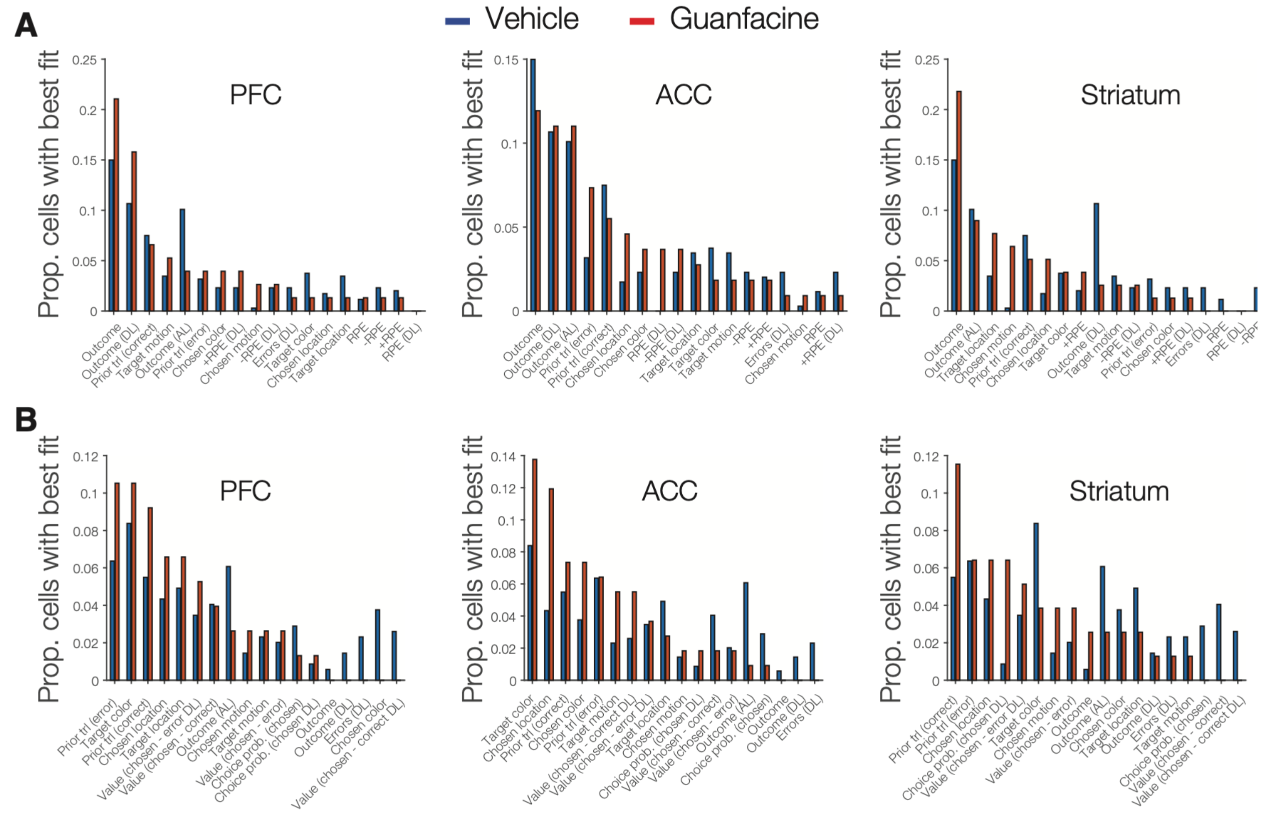


**Figure S5.** The best fit regressors for explaining the activity of each respective brain region. (**A**) The number of neurons whose activity during the feedback epoch each is best accounted for by the regressor denoted on the x-axis - ranked in descending order for neurons in the Guanfacine (*red*) and vehicle (*blue*) condition. The similarity in rankings were tested for via Kendall's tau correlation. From left to right: the dlPFC ($p = .006$; Tau = .477), the ACC ($p = .014$; Tau = .425), and the striatum ($p = .014$; Tau = .425). (**B**) Same format as *A* for the attention epoch. From left to right: the dlPFC ($p = .970$ n.s.; Tau = -.007), the ACC ($p = .112$ n.s.; Tau = -.281), and the striatum ($p = .175$ n.s.; Tau = .242).

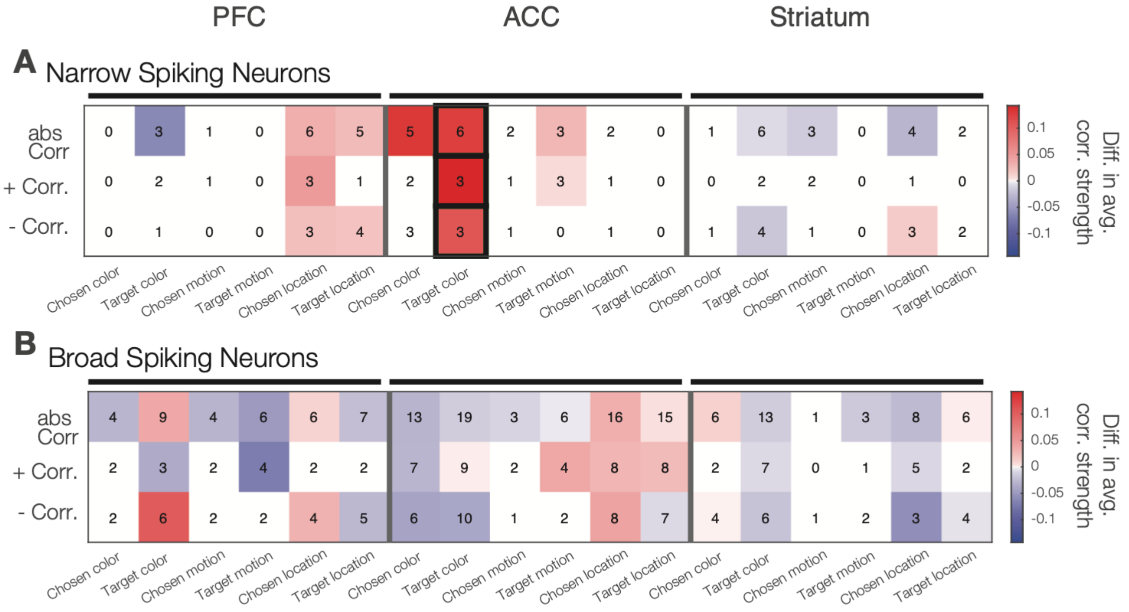


**Figure S6.** Guanfacine enhanced correlations of narrow spiking neurons firing with the target color during the attention epoch. (**A**) The difference in average correlation strength in the Guanfacine versus vehicle condition for narrow spiking neurons for variables characterizing the target stimulus and the chosen stimulus. Thick outlined squares denote significant differences at $p<0.05$. (**B**) Same format as *A* for broad spiking neurons.

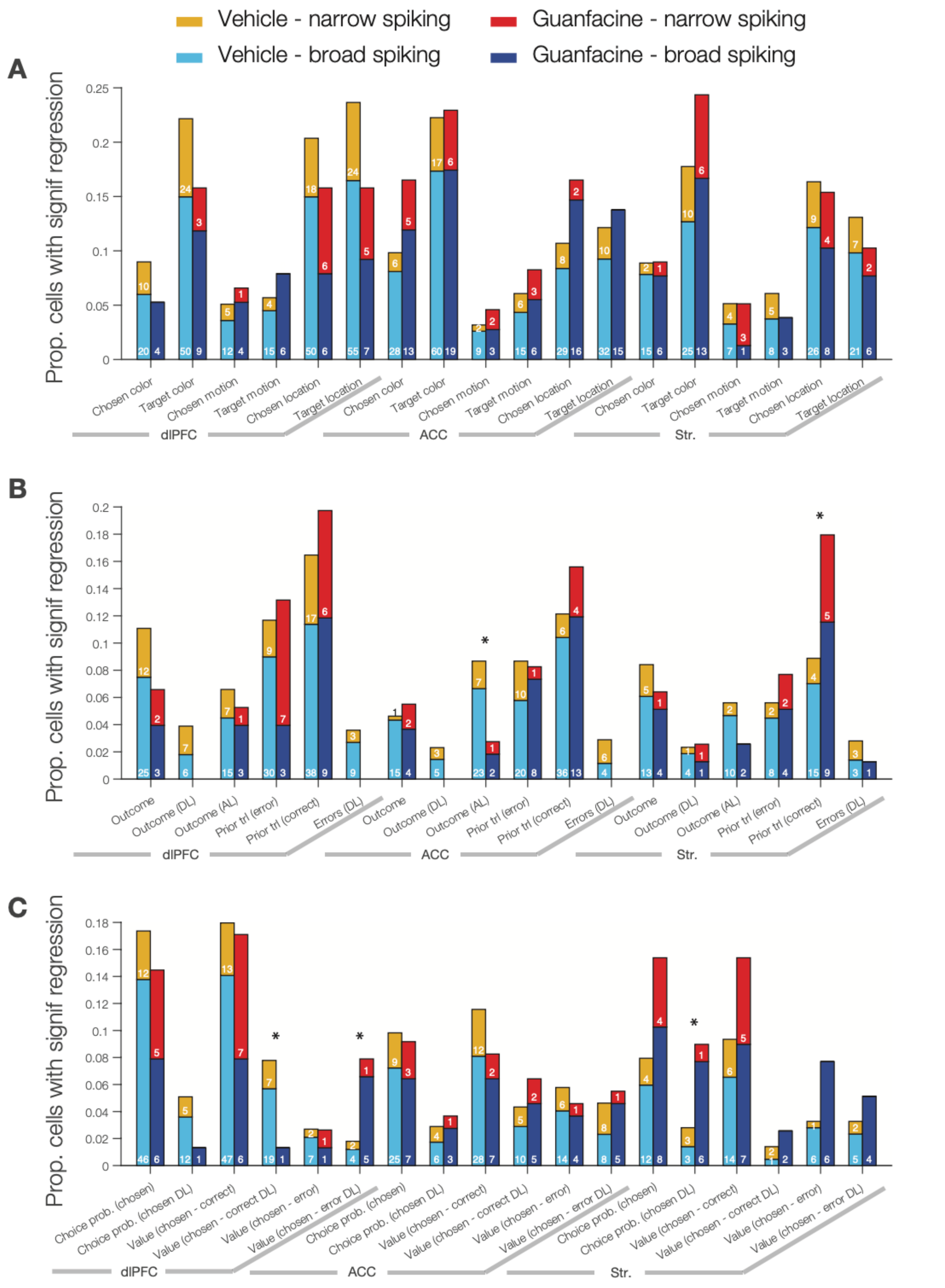


**Figure S7.** The proportion of neurons with firing during the attention epoch that significantly regress to target stimulus variables (color, direction of motion, location) (*A*), outcome variables (*B*) and model-derived reward prediction errors (*C*). A star above a bar indicates a significant difference of Guanfacine and vehicle conditions.
